# Mendelian Randomization evaluation of causal effects of fibrinogen on incident coronary heart disease

**DOI:** 10.1101/448381

**Authors:** Cavin K. Ward-Caviness, Paul S. de Vries, Kerri L. Wiggins, MS, Jennifer E. Huffman, Jennifer E. Huffman, Lisa R. Yanek, Lawrence F. Bielak, Franco Giulianini, Xiuqing Guo, Marcus E. Kleber, Tim Kacprowski, Stefan Groβ, Astrid Petersman, George Davey Smith, Fernando P. Hartwig, Jack Bowden, Gibran Hemani, Martina Müller-Nuraysid, Konstantin Strauch, Wolfgang Koenig, Melanie Waldenberger, Thomas Meitinger, Nathan Pankratz, Eric Boerwinkle, Weihong Tang, Yi-Ping Fu, Andrew D Johnson, Ci Song, Ci Song, Moniek P.M. de Maat, André G. Uitterlinden, Oscar H. Franco, Jennifer A. Brody, Barbara McKnight, Yii-Der Ida Chen, Bruce M. Psaty, Rasika A. Mathias, Diane M. Becker, Patricia A. Peyser, Jennifer A. Smith, Suzette J. Bielinski, Paul M. Ridker, Kent D. Taylor, Jie Yao, Russell Tracy, Graciela Delgado, Stella Trompet, Naveed Sattar, J. Wouter Jukema, Lewis C. Becker, Sharon L.R. Kardia, Jerome I. Rotter, Winfried März, Marcus Dörr, Daniel I. Chasman, Abbas Dehghan, Christopher J. O’Donnell, Nicholas L. Smith, Annette Peters, Alanna C. Morrison

## Abstract

**Background:** Fibrinogen is an essential hemostatic factor and cardiovascular disease risk factor. Early attempts at evaluating the causal effect of fibrinogen on coronary heart disease (CHD) and myocardial infraction (MI) using Mendelian randomization (MR) used single variant approaches, and did not take advantage of recent genome-wide association studies (GWAS) or multi-variant, pleiotropy robust MR methodologies.

**Methods and Findings:** We evaluated evidence for a causal effect of fibrinogen on both CHD and MI using MR. We used both an allele score approach and pleiotropy robust MR models. The allele score was composed of 38 fibrinogen-associated variants from recent GWAS. Initial analyses using the allele score incorporated data from 11 European-ancestry prospective cohorts to examine incidence CHD and MI. We also applied 2 sample MR methods with data from a prevalent CHD and MI GWAS. Results are given in terms of the hazard ratio (HR) or odds ratio (OR), depending on the study design, and associated 95% confidence interval (CI).

In single variant analyses no causal effect of fibrinogen on CHD or MI was observed. In multi-variant analyses using incidence CHD cases and the allele score approach, the estimated causal effect (HR) of a 1 g/L higher fibrinogen concentration was 1.62 (CI = 1.12, 2.36) when using incident cases and the allele score approach. In 2 sample MR analyses that accounted for pleiotropy, the causal estimate (OR) was reduced to 1.18 (CI = 0.98, 1.42) and 1.09 (CI = 0.89, 1.33) in the 2 most precise (smallest CI) models, out of 4 models evaluated. In the 2 sample MR analyses for MI, there was only very weak evidence of a causal effect in only 1 out of 4 models.

**Conclusions:** A small causal effect of fibrinogen on CHD is observed using multi-variant MR approaches which account for pleiotropy, but not single variant MR approaches. Taken together, results indicate that even with large sample sizes and multi-variant approaches MR analyses still cannot exclude the null when estimating the causal effect of fibrinogen on CHD, but that any potential causal effect is likely to be much smaller than observed in epidemiological studies.

*Author Summary:* Initial Mendelian Randomization (MR) analyses of the causal effect of fibrinogen on coronary heart disease (CHD) utilized single variants and did not take advantage of modern, multivariant approaches. This manuscript provides an important update to these initial analyses by incorporating larger sample sizes and employing multiple, modern multi-variant MR approaches to account for pleiotropy. We used incident cases to perform a MR study of the causal effect of fibrinogen on incident CHD and the nested outcome of myocardial infarction (MI) using an allele score approach. Then using data from a case-control genome-wide association study for CHD and MI we performed two sample MR analyses with multiple, pleiotropy robust approaches. Overall, the results indicated that associations between fibrinogen and CHD in observational studies are likely upwardly biased from any underlying causal effect. Single variant MR approaches show little evidence of a causal effect of fibrinogen on CHD or MI. Multi-variant MR analyses of fibrinogen on CHD indicate there may be a small positive effect, however this result needs to be interpreted carefully as the 95% confidence intervals were still consistent with a null effect. Multi-variant MR approaches did not suggest evidence of even a small causal effect of fibrinogen on MI.

## Introduction

Fibrinogen is an essential component of the clotting and hemostasis system with a strong genetic basis [1-3]. Although it primarily serves as the precursor to fibrin, it also carries out several other functions, including enhancing platelet aggregation and mediating inflammation [4, 5]. In epidemiologic studies, fibrinogen levels are associated with coronary heart disease (CHD) [6-8], myocardial infarction (MI) [9, 10], ischemic stroke [11, 12], and abdominal aortic aneurysm [13, 14].

Mendelian randomization (MR) is an instrumental variable analysis method which uses genetic variants as instruments to uncover evidence for a causal relationship between a modifiable risk factor and outcome.[15] MR studies utilizing a limited number of genetic variants in the *FGB* promoter have yielded little evidence of a causal effect of fibrinogen on CHD or MI [16-18]. In a genome-wide association study (GWAS) for fibrinogen, each fibrinogen-associated variant was individually evaluated for association with CHD, but no associations provided substantial evidence of a causal effect [19]. To date MR studies of fibrinogen have been limited to single variant approaches which have not taken into account recent GWAS findings or modern, multi-variant MR methodologies. Here we re-examine the potential for fibrinogen to be a causal biomarker for CHD and MI, taking into account these improved approaches.

## Results

For incident CHD there were 3,147 incident events observed in 15,427 participants in the discovery analyses, and 1,482 incident events among the 34,209 participants in the replication analyses. Of the 18,798 participants in the incident MI discovery analyses, 1,711 had an incident MI. For the replication analyses, there were 687 incident MI events out of the 33,288 participants. **Table 1** contains the distributions of clinical covariates and fibrinogen. The FGB variant rs1800790 (commonly used in previous fibrinogen MR analyses) had a weaker association (by effect size) than the allele score (**Supplemental Table 3**). In single variant analyses of rs1800790 the estimated causal effect appeared to be centered around the null with little evidence of a causal effect of fibrinogen on CHD or MI (**Supplemental Table 3**), consistent with published literature. In multi-variant MR using the 2SC model, we observed evidence of a causal association of fibrinogen on incident CHD in the discovery and replication analyses which remained in a combined analysis of all cohorts (HR = 1.75; CI = 1.22-2.51; P = 0.002; **Figure 2**). For incident MI, we observed an elevated HR that included the null, even in the combined analysis (HR = 1.45; CI = 0.85-2.49; P = 0.17; **Figure 3**). ***Pleiotropy robust models***

**Table 1.** Clinical Covariates. Clinical covariates for all participating cohorts. KORA did not have incident CHD data and thus did not participate in these analyses. GENOA and LURIC had too few incident MI cases for analysis. * For FHS 172 individuals were not current smokers but were not distinguished as former vs never smokers thus percentages were not computed for these categories and the N for those with information is given. ** For SHIP only interval censored data was available. Follow-up time represents the time from initial exam to final exam. BMI = body mass index; CHD = coronary heart disease; HDL = high-density lipoprotein cholesterol; LDL = low-density lipoprotein cholesterol; MI = myocardial infarction; NA = not available

|  | Discovery |  |  |  |  |  |  | Replication |  |  |  |
| --- | --- | --- | --- | --- | --- | --- | --- | --- | --- | --- | --- |
|  | KORA | GENOA | ARIC | CHS | GeneSTAR | FHS | RS | MESA | WGHS | LURIC | SHIP |
| **N** | 3,788 | 417 | 8,815 | 2,939 | 594 | 1,828 | 834 | 2,506 | 27,553 | 921 | 3,229 |
| **Mean (SD)** |  |  |  |  |  |  |  |  |  |  |  |
| MI follow-up time (y) | 8.44 (1.52) | - | 20.8 (6.03) | 10.3 (6.0) | 6.72 (3.59) | 17.9 (5.32) | 9.32 (3.16) | 6.25 (1.12) | 17.5 (3.76) | - | 11.14 (0.8)** |
| CHD follow-up time (y) | - | 10.4 (3.6) | 20.0 (6.49) | 10.3 (5.9) | 6.07 (3.19) | 17.9 (5.32) | 9.18 (3.24) | 6.18 (1.23) | 17.4 (3.95) | 9.43 (2.55) | 11.14 (0.8)** |
| Age (y) | 49.2 (13.9) | 58.6 (10.0) | 54.2 (5.68) | 72.4 (5.4) | 51.1 (11.3) | 53.8 (10.0) | 72.2 (6.83) | 62.7 (10.2) | 54.6 (7.08) | 60.1 (11.5) | 46.39 (15.11) |
| BMI (kg/m<sup>2</sup>) | 27.2 (4.69) | 30.6 (6.3) | 27.0 (4.86) | 26.2 (4.4) | 29.3 (6.3) | 27.4 (4.93) | 26.7 (3.80) | 27.7 (5.10) | 25.1 (6.78) | 27.5 (4.36) | 27.0 (4.75) |
| Fibrinogen (g/L) | 2.63 (0.60) | 3.19 (0.80) | 2.95 (0.61) | 3.14 (0.61) | 3.83 (1.08) | 3.14 (0.61) | 3.95 (0.87) | 3.35 (0.70) | 3.59 (0.78) | 3.69 (0.90) | 2.95 (0.68) |
| log(Fibrinogen) | 0.94 (0.22) | 1.11 (0.36) | 3.97 (0.11) | 1.13 (0.19) | 1.30 (0.27) | 1.12 (0.19) | 1.35 (0.21) | 1.19 (0.20) | 1.25 (0.22) | 1.28 (0.26) | 1.06 (0.22) |
| HDL (mg/dL) | 57.9 (17.0) | 52.5 (14.3) | 51.1 (16.7) | 55.8 (15.9) | 52.1 (15.8) | 51.3 (15.3) | 204 (24.5) | 52.4 (15.7) | 53.1 (16.6) | 41.8 (11.7) | 57.04 (17.1) |
| LDL (mg/dL) | 137 (41.4) | 117 (31.0) | 137 (37.6) | 134 (35.8) | 130 (41.6) | 126 (33.0) | 104 (12.5) | 117 (30.3) | 122 (37.3) | 121 (33.7) | 137.3 (44.2) |
| **N (%)** |  |  |  |  |  |  |  |  |  |  |  |
| MI | 109 (2.88) | - | 859 (9.74) | 498 (16.9) | 30 (5.05) | 158 (8.64) | 57 (6.83) | 62 (2.50) | 413 (1.50) | - | 212 (6.5) |
| CHD | - | 77 (18.5) | 1,625 (18.4) | 942 (32.1) | 85 (14.31) | 305 (16.7) | 113 (13.6) | 128 (5.10) | 1,035 (3.76) | 59 (6.41) | 260 (8.0) |
| Sex (male) | 1,854 (48.9) | 156 (37.4) | 3,983 (45.2) | 1,792 (61.0) | 321 (54.0) | 829 (45.0) | 421 (50.5) | 1,308 (52.2) | 0 (0.00) | 499 (54.2) | 1,537 (47.6) |
| Current Smokers | 962 (25.4) | 47 (11.3) | 2,633 (29.9) | 322 (11.0) | 135 (22.8) | 341 (18.7) | 144 (17.3) | 286 (11.4) | 3200 (11.6) | 197 (21.4) | 1,096 (33.9) |
| Former Smokers | 1,262 (33.3) | 157 (37.6) | 2,050 (23.3) | 1,216 (41.4) | 202 (34.0) | 404* | 429 (51.4) | 1,109 (44.4) | 10,096 (36.6) | 273 (29.6) | 1,004 (31.1) |
| Never Smokers | 1,560 (41.2) | 213 (51.0) | 4,109 (46.6) | 1401 (47.7) | 257 (43.3) | 911* | 261 (31.3) | 1,104 (44.2) | 14,233 (51.7) | 451 (49.0) | 1,129 (35.0) |
| Hypertension | 1,079 (28.5) | 310 (74.3) | 2,272 (25.8) | 1549 (52.7) | 261 (43.94) | 563 (30.8) | 204 (24.5) | 1,097 (43.8) | 6,654 (24.2) | 625 (67.5) | 671 (20.8) |
| Type 2 Diabetes | 565 (14.9) | 52 (12.5) | 1,659 (18.8) | 349 (11.9) | 59 (9.93) | 129 (7.10) | 104 (12.5) | 150 (6.70) | 0 (0.00) | 283 (30.7) | 174 (5.4) |

**Figure.1.**
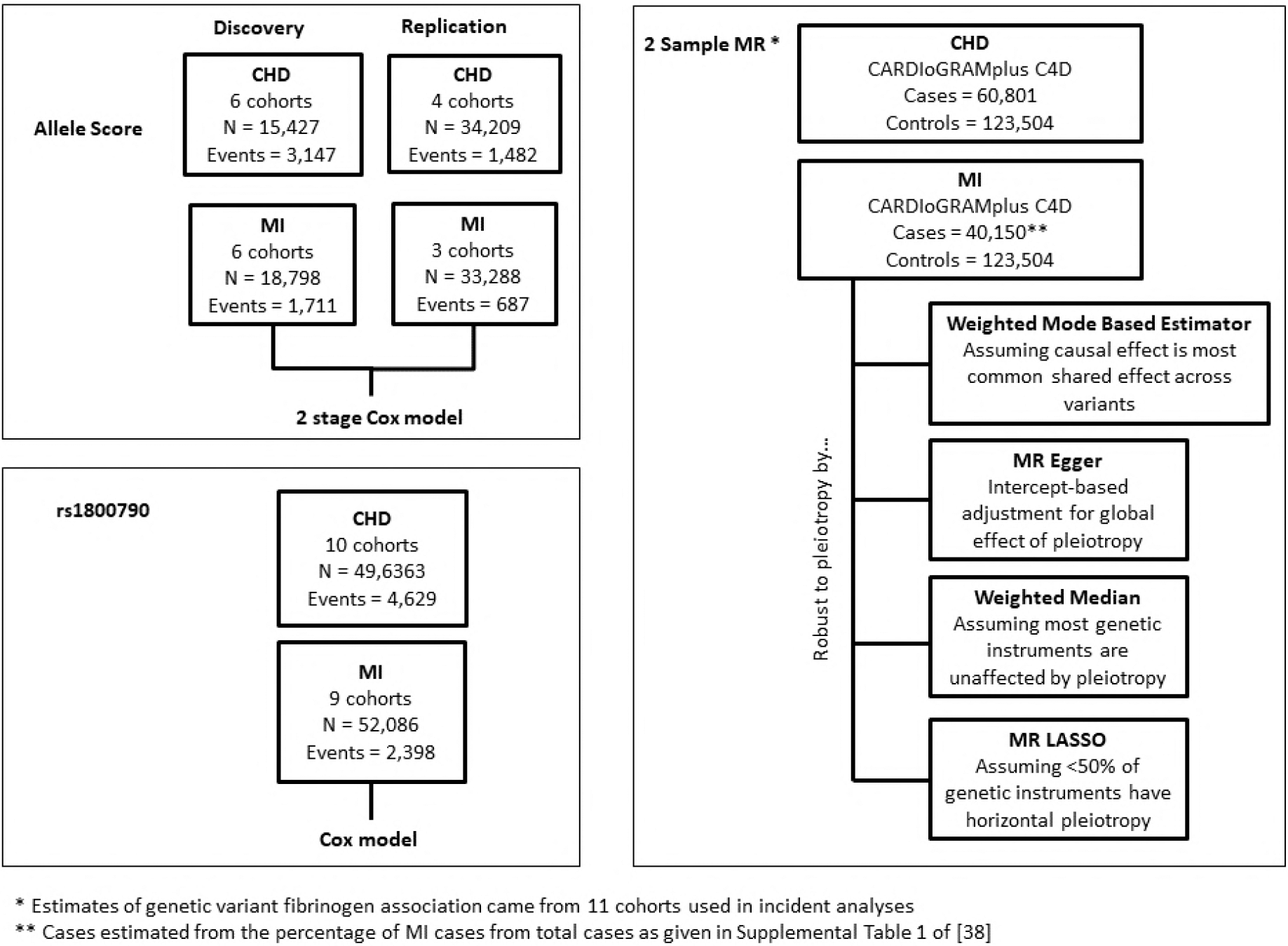
Study Outline. Outline of analyses using the allele score, rs1800790 and 2 Sample MR approaches including the analytic method used to estimate the causal effect, subject to valid MR assumptions, for all stages of the analysis. CHD = coronary heart disease; MR = Mendelian Randomization; MI = myocardial infarction

**Figure.2.**
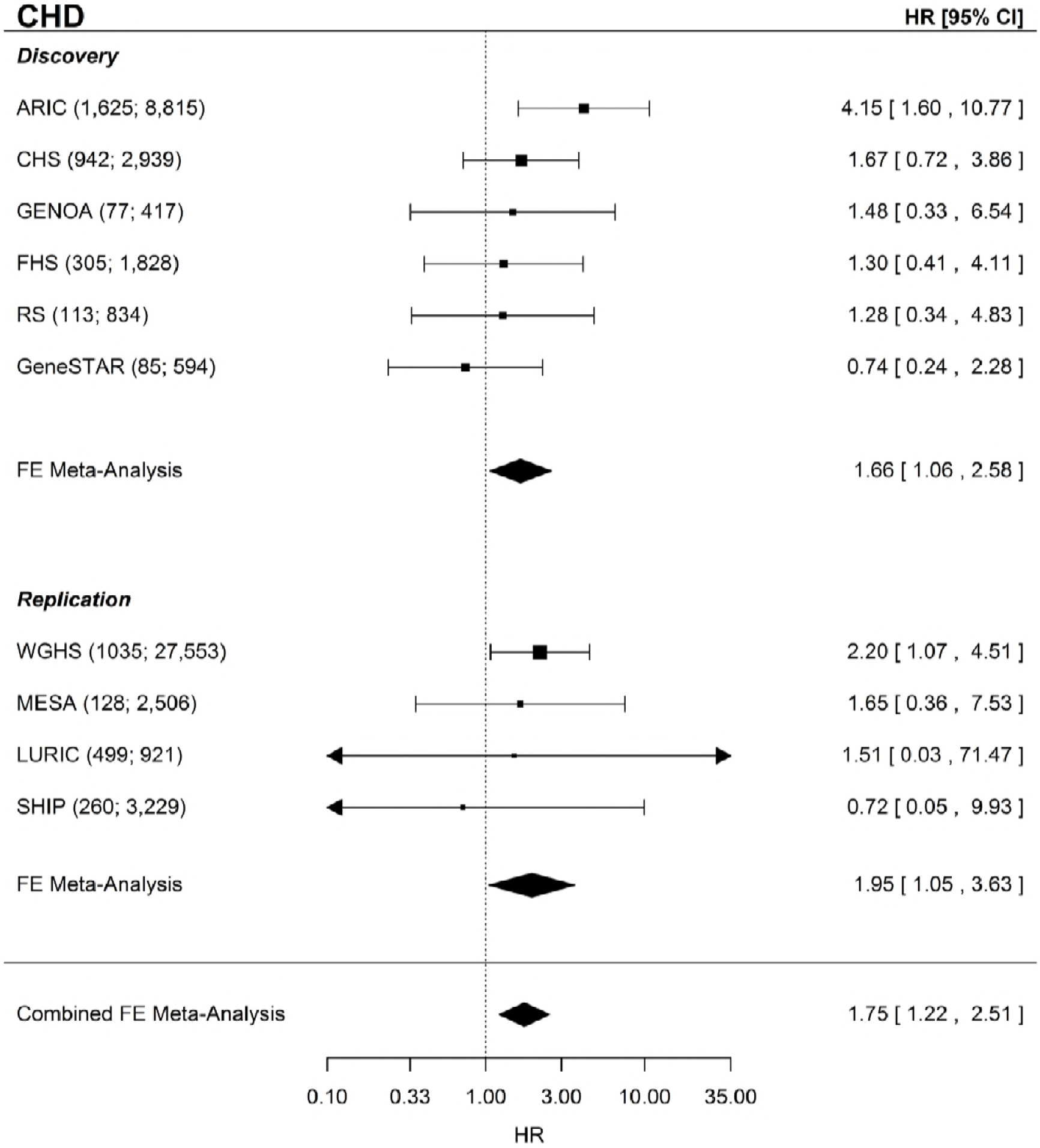
CHD Forest Plot. Forest plot of the CHD MR analysis for the discovery, replication, and combined sets of cohorts. Shown beside each cohort name is the sample size and number of incident CHD events given as (N events; N total). CHD = coronary heart disease; FE = fixed-effects; HR = hazard ratio; CI = confidence interval

**Figure.3.**
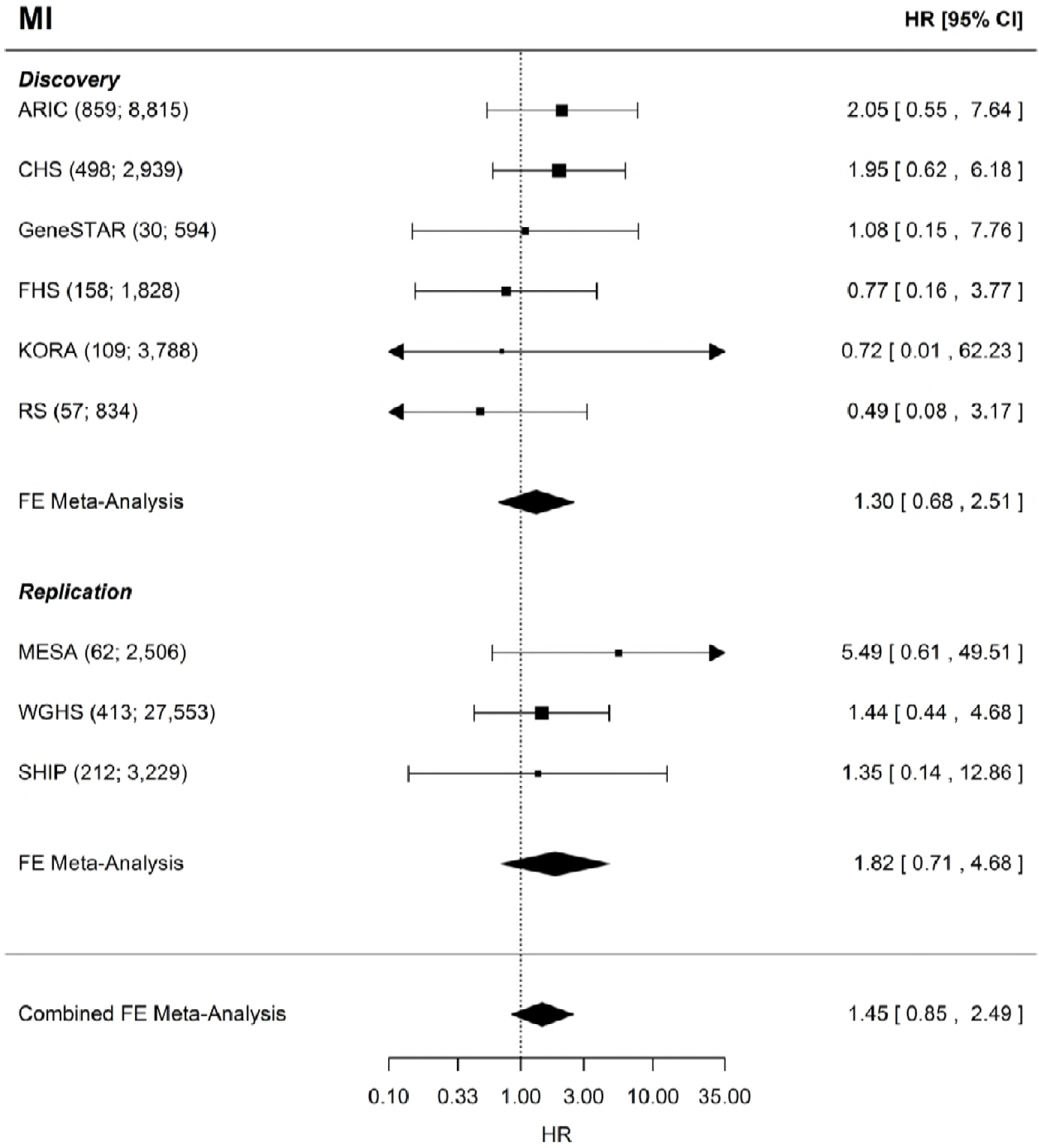
MI Forest Plot. Forest plot of the MI MR analysis for the discovery, replication, and combined sets of cohorts. Shown beside each cohort name is the sample size and number of incident MI events given as (N events; N total). MI = myocardial infarction, FE = fixed-effects, HR = hazard ratio, CI = confidence interval

In sensitivity analyses four MR methods were used each of which is at least partially robust to horizontal pleiotropy under differing assumptions. For CHD, three of the four models showed a positive effect, albeit smaller than the effect observed in the 2SC model, with the MR PRESSO method having the largest causal OR (OR = 1.18; CI = 0.98, 1.42; **Table 2**). For MI only the MR PRESSO method showed a causal OR > 1 (OR = 1.16; CI = 0.98, 1.38; **Table 2**), again substantially reduced from that observed in the 2SC model. All other models for MI showed little evidence of a causal effect of fibrinogen on MI.

**Table 2.** Multi-variant, Pleiotropy Robust MR Methods. To further examine potential effects of pleiotropy we ran several multi-variant, pleiotropy robust models including MR Egger, Weighted Mode Based Estimator (MBE), Weighted Median, and MR PRESSO. Each uses a different means to account for pleiotropy and has different assumptions used to estimate the causal effect in the presence of pleiotropy. Odds ratios are per 1 g/L increase in genetically determined fibrinogen. CHD = coronary heart disease; CI = confidence interval; MI = myocardial infarction; MR = mendelian randomization

| Method | Robust to pleiotropy by ... | CHD Causal OR (95% CI) | MI Causal OR (95% CI) |
| --- | --- | --- | --- |
| MR Egger | Intercept-based adjustment for global effect of pleiotropy | 0.98 (0.70, 1.39) | 0.89 (0.63, 1.26) |
| Weighted MBE ($\phi = 1$) | Assuming causal effect is most common shared effect across variants | 1.09 (0.89, 1.33) | 0.98 (0.79, 1.21) |
| Weighted Median | Assuming most ($\geq 50\%$) genetic instruments are unaffected by pleiotropy | 1.12 (0.91, 1.37) | 1.03 (0.82, 1.29) |
| MR PRESSO | Assuming $<50\%$ of genetic instruments have horizontal pleiotropy | 1.18 (0.98, 1.42) | 1.17 (0.98, 1.40) |

As a further test we examined MR associations of fibrinogen on CHD risk using published data available in the MR-Base. While some of the CHD risk factors showed a positive causal effect estimate, none provided substantial evidence for excluding the null after accounting for the number of tests performed (**Supplemental Table 5**).

## Discussion

The attractiveness of fibrinogen as a causal factor in CHD comes from its roles in both thrombosis and inflammation. Fibrinogen is the precursor to fibrin, which interlinks into a mesh that acts as the scaffold of blood clots. Additionally, fibrinogen also has an active role in platelet aggregation,[20] thus contributing to the formation of platelet plugs. By binding the CD11b/CD18 integrin receptor fibrinogen activates the NF-κB pathway [5], an important pathway in inflammation as well as the formation, destabilization, and rupture of atherosclerotic plaques [21, 22]. As a modifiable risk factor [23] even a small causal effect of fibrinogen on CHD could have substantial public health implications.

Using the allele score approach, a 1 g/L higher fibrinogen concentration was causally associated with a HR of 1.75 (CI = 1.22-2.51) in the combined cohort analysis for CHD. However, sensitivity analysis using methods robust to pleiotropy arising from independent effects of SNPs on exposure and outcome (which could invalidate MR analyses) suggested a substantially weaker causal effect on CHD even for the model with the strongest effect estimate (OR = 1.18 per 1 g/L higher fibrinogen; CI = 0.98, 1.42), and the MR Egger model showed virtually no evidence of a causal effect-though the wide 95% confidence interval encompassed effects from all other models. Overall, when accounting for potential horizontal pleiotropy, the accumulated evidence points to a substantially weaker casual effect of fibrinogen on CHD than the observational risk ratio of 1.8 (CI = 1.6, 2.0) previously reported [6]. Using rs1800790 in a single variant MR analysis, there was limited evidence of any causal effect, though the 95% confidence interval could not exclude positive estimated causal effects seen in multi-variant analyses. In combination these analyses suggest that when after accounting for horizontal pleiotropy the effect of fibrinogen on CHD is likely to be small and that current MR estimates of the potential causal effect remain unable to exclude the null despite large sample sizes and the latest methodologies.

### Comparison with previous MR analyses

Previous MR studies assessing the causal effect of fibrinogen on CHD or MI focused exclusively on rs1800790 [24, 25]. In a few studies one additional variant also in the *FGB*promoter region was examined, however this variant is in nearly complete LD with rs1800790, particularly in Europeans [16, 18]. The allele score was a better predictor of fibrinogen than rs1800790 alone (**Supplemental Table 3**). Though the allele score estimated a causal effect of fibrinogen on CHD similar to observational studies, much of this appeared to be driven by pleiotropy as estimated effects decreased in models more robust to pleiotropy (**Table 2**). This highlights the need to balance increased power from multi-variant approaches with the potential for increased pleiotropy in these instruments.

For the CHARGE cohorts we used exclusively incident cases whereas previous studies utilized populations composed entirely or primarily of prevalent cases. In some instances, the use of prevalent cases may bias MR studies such as if the disease subsequently what is perceived as a disease risk factor, e.g. if CHD leads to higher fibrinogen as opposed to the reverse, then reverse confounding can still occur even in an MR setting [26]. Additionally, if the risk factor were to affect severity of an event, e.g. the fatality of MI, then use of prevalent cases may dilute the MR-estimated causal effect as the most severe cases may not be observed due to being too ill to participate or suffering a fatal event. This type of prevalence-incidence bias is not exclusive to MR analyses [27-29]. However, care must still be taken when interpreting results from incident case MR studies as the exclusion of prevalent cases is equivalent to conditioning on disease status at baseline. This has the potential to introduce bias in the form of an exclusion restriction violation.[30] Whether bias is introduced and the degree of confounding are dependent on the actual biological processes that account for the relationship between the genetic instrument(s) chosen, the modifiable risk factor, and outcome in the MR analysis. When performing incident case MR it is best to combine the efforts with

MR analyses including prevalent cases and interpret results for both with careful consideration towards their underlying assumptions, strengths, and weaknesses. In general, our results are compatible with previous MR studies, however we use more modern methods, including multi-variant, pleiotropy robust methods, able to produce smaller confidence intervals and which indicate that after accounting for pleiotropy there may be a small positive effect of fibrinogen on CHD. This is particularly true for the methods producing the most precise estimates. However, these results warrant further investigations as confidence intervals for some models were still wide and with results for the single variant and MR Egger analyses possibly more consistent with no causal effect than even a small causal effect.

### Strengths and limitations

As with all MR studies the causal effects estimated here are based on regression estimates for genetic variants and are only valid, causal estimates under the assumptions of MR. Additionally, causal estimates generated via MR methodologies are for lifelong, genetically determined increases in the exposure, e.g. fibrinogen, which means that caution should be exercised when applying clinical interpretations or attempting to translate results into estimates of an intervention.[31, 32]This study had some overlap between studies involved in the GWAS used to select fibrinogen variants and those used in the MR analyses. Our approach to mitigate this was to replicate the allele score analysis in an independent set of cohorts. For the pleiotropy robust 2-sample MR approaches this overlap was unavoidable, however there was no overlap for the cases which means that unbiased estimates should be obtained [33]. A strength of the study is the use of incident cases for the allele score model approach which reduces the potential for bias from reverse confounding (which can still affect MR studies) and prevalence-incidence bias. Additionally, even though the allele score approach was sensitive to horizontal pleiotropy we used an array of additional approaches that were each partially robust to horizontal pleiotropy through different assumptions about the nature of the pleiotropy. These models often have lower power than other approaches, which motivated our use of a previously published GWAS which had 60,801 prevalent cases and 123,504 controls [34]. However, to prevent potential bias and more closely align with our initial analyses, a large sample size of incident cases independent of those used to evaluate associations between genetic variants and fibrinogen would have been preferable.

## Conclusion

Fibrinogen represents an important role in thrombosis, platelet aggregation, and inflammation making it a promising risk factor for CHD. Despite the epidemiological evidence, MR studies using prevalent cases and single variant approaches have consistently shown no causal effect of fibrinogen on CHD. Out results indicate that epidemiologic studies may substantially overestimate any causal effect of fibrinogen on CHD. While some MR models which accounted for pleiotropy did show a modest causal effect, the 95% confidence intervals still contained the null indicating that researchers should exercise caution in interpreting these results. Further analyses using larger sample sizes and more precise methods are warranted to better resolve the effect of fibrinogen on CHD.

## Methods

This study was conducted within the Cohorts for Heart and Aging Research in Genomic Epidemiology (CHARGE) consortium [35] using 11 European-ancestry cohorts. For incident CHD, six cohorts participated in the initial (discovery) analyses (N = 15,427), and four cohorts (N = 34,209) contributed data for replication. For incident MI, six cohorts participated in the discovery (N = 18,798), and three cohorts participated in the replication (N = 33,288) analyses. Details on all cohorts are given in the Supplemental Materials and the clinical covariates in **Table 1**, and **Figure 1** outlines all analyses. Data collection analysis for all cohorts was approved by their respective Institutional Review Boards and/or ethical committees, and all cohorts obtained written, informed consent from participants.

### Assessment of CHD and MI

We defined incident CHD as validated, incident fatal or non-fatal CHD events which included: validated hospitalized MI, CHD-related hospitalizations, definite CHD deaths, likely CHD deaths, and CHD-related revascularization procedures, e.g. percutaneous coronary intervention and coronary artery bypass grafting. Incident MI was defined as a validated fatal or non-fatal MI and included definite MI hospitalizations. For cohorts that used questionnaires as a component of the follow-up procedures, all events were corroborated with medical records and/or review by trained medical personnel. Cohort specific details are given in the Supplemental Online Methods.

### Fibrinogen Assessment

Fibrinogen was assessed by a variety of methods, with seven cohorts using the Clauss method [36]. Of the remaining four cohorts, RS used a clotting time-derived method to assess fibrinogen concentrations, while KORA, MESA, and WGHS used immunological assays to assess total fibrinogen.

### Genotyping and Imputation

Genotyping and imputation were performed separately in all cohorts, per published methods [37]. All participating studies used either the HapMap build 36 [38], 1000 Genomes phase I version 3, or 1000 Genomes phase I version 2 reference panel for imputation [39]. Imputation was performed via MACH[40] or IMPUTE [41]. Low quality variants were excluded in line with previously published approaches: MACH imputation quality < 0.3 or IMPUTE imputation quality < 0.4 [37].

### Creation of the Allele Score

We evaluated 69 variants associated with fibrinogen in at least one of three recent genome/exome-wide association studies [2, 19, 37] for inclusion into the allele score [42]. We applied four criteria to each variant to improve the plausibility that each meets the MR assumptions. First, to ensure that the variants were not correlated with known risk factors for cardiovascular disease (CVD), the Spearman correlation between each of the variants and body mass index (BMI), low-density lipoprotein (LDL) cholesterol, high-density lipoprotein (HDL) cholesterol, type 2 diabetes mellitus (binary), hypertension (binary), and smoking (ever, never, current) was tested within each cohort and any variants with a Spearman correlation greater than 0.10 in any cohort for any of these outcomes were removed. Second, the variants were tested for linkage disequilibrium (LD) with known CHD loci [34, 43-52] using SNAP from the Broad Institute with LD patterns coming from European ancestry individuals [53]. As no variant had r^2^ > 0.20 with a CHD locus, they were considered independent of known CHD loci. Next, we reduced pairs of variants in high LD (r^2^ > 0.70) by preferentially retaining those variants that were found in the largest genome-wide scan [37]. Finally, we eliminated any variants that were missing across any of the discovery cohorts, leaving 38 variants that composed the allele score (**Supplemental Table 1**). We tested the allele score for association with each of the aforementioned CHD risk factors in each cohort as well as in a meta-analysis of all cohorts. The allele score was not associated with any CHD risk factor in the meta-analysis after a Bonferroni correction for the six tests performed (P > 0. 008; **Supplemental Table 2**). Six variants from the allele score which were unavailable in one or more replication cohorts were removed from the allele score in the replication phase to ensure a consistent allele score in the replication meta-analysis (**Supplemental Table 1**). In a sensitivity analysis these variants were also removed from the discovery cohorts and the causal effect evaluated in a combined meta-analysis.

Each genotype was aligned prior to summing to create the score so that the designated effect allele corresponded to a positive association with fibrinogen according to the direction of effect in the largest and most recent fibrinogen GWAS [19].

### Mendelian Randomization

MR is a powerful framework that uses genetic variants as instrumental variables to infer causal relationships between a defined exposure and outcome. The causal effect estimated by MR is the alteration in exposure due to genetic variation and is thus assumed to be over the entire life course. There are three assumptions for a genetic variant to be a valid instrument for MR[54]

1. The genetic variant is independent of confounders of exposure and outcome under examination
2. The genetic variant is associated with the exposure
3. The genetic variant is independent of the outcome conditional on the exposure and any confounders

In addition to these three conditions, valid estimates from MR are dependent on any parametric assumptions of the model being used to estimate relevant coefficients and standard errors.

Our initial MR analyses used a two-stage procedure employing a Cox regression model (2SC). To improve power, we regressed fibrinogen on age and sex and used the resulting residuals as input to the 2SC analyses. In the first stage of the 2SC procedure the fibrinogen residuals were regressed on the allele score. In the second stage the predicted values from the first stage regression were associated with incident MI or CHD via a Cox proportional hazards model. This approach is similar to the two-stage predictor-substitution MR approach [55-57], and results from the 2SC model are given per unit (g/L) increase in the fibrinogen residuals. We used a fixed effects model for all meta-analyses since we observed little heterogeneity according to the Q-statistics [58] (P(Q) > 0.05 for all analyses). We also compared associations with our allele score to those obtained using a single variant, FGB-455G>A (rs1800790), which is a commonly used variant for fibrinogen MR analyses [16, 18]. We performed sensitivity analyses using four pleiotropy robust methods each of which uses a different approach to partially relax the no horizontal pleiotropy assumption of MR analyses:[59] MR-Egger [54], MR mode based estimate (MBE),[60] MR PRESSO,[61] and Weighted median [62]. For these sensitivity analyses, we used the prevalent CHD and MI GWAS results from CARDIoGRAMplusC4D consortium [34] as it had a larger sample size (60,801 prevalent cases and 123,504 controls) and these methods often have lower power to detect effects. For estimates of variant effects on fibrinogen we used fixed-effects met-aanalysis estimates from the 11 cohorts in these analyses. Since an individual cannot be both a prevalent and incident CHD or MI case at the same sampling, there was no overlap amongst the cases between our incident analyses and the prevalent cases used in the CARDIoGRAMplusC4D GWAS. There would still be some overlap amongst the non-cases/controls which could bias estimates towards the null.

We also examined whether fibrinogen showed evidence for a causal effect on 7 metabolic CHD risk factors using MR-base (www.mrbase.org), a database of published GWAS available for MR [63]. We focused on metabolic CHD risk factors as initial results indicated that body mass index was the trait with which our allele score showed the strongest evidence for pleiotropy - potentially horizontal (i.e. SNPs affecting fibrinogen and CHD via independent pathways) and vertical (i.e. fibrinogen-associated SNPs also associated with risk factors downstream of fibrinogen) as the associations did not distinguish between the two.

The CHD risk factors were body mass index [64], waist circumference [65], waist-to-hip ratio [65], low-density lipoprotein cholesterol,[66] triglycerides [66], homeostatic model assessment insulin resistance (HOMA-IR) [67], and Type 2 diabetes [68]. As MR-base only contains published GWAS we used the most recently published GWAS for fibrinogen for our variant-fibrinogen associations [1] but limited to those variants present in our allele score. For the CHD risk factors we compared causal effect estimates obtained from the inverse variance weighted method (which assumes no unbalanced horizontal pleiotropy), to those from the pleiotropy robust MR Egger, and Weighted median methods. All three methodologies were implemented in MR-base.

Statistical analyses were performed in R.[69] Meta-analyses were performed using the R package *metafor.[70]* Cox models were estimated via the *coxph* function in the R package survival[71] with the exception of SHIP where the *survreg* function was used with an exponential distribution to account for the interval censored data. MR-Egger and weighted median results were performed using the R package *MendelianRandomization* and *Two Sample* MR.[63] MR MBE analyses were performed using the methods given by Hartwig *et*al.[60] The default bandwidth (φ = 1) was used for MR MBE as results did not show sensitivity to the choice of bandwidth. MR PRESSO analyses were performed using code available at the MR PRESSO GitHub repository (https://github.com/rondolab/MR-PRESSO)[61]. We used the robust MR estimates from MR PRESSO which are equivalent to performing an inverse-variance weighted MR analysis after removing outlying variants, which may be influenced by horizontal pleiotropy, as identified by MR PRESSO. Results from the 2SC model are reported in terms of the hazard ratio (HR), while all results that utilize the prevalent disease GWAS are reported in terms of the odds ratio (OR). All HR and OR are given per 1 g/L higher fibrinogen. All confidence intervals (CI) reported are 95% CI.

## Author Contributions

JIR, CJO’D, MD, DMB, LCB, JWK, CKW-C, PMR, DIC, SLRK, NLS, A Peters, ACM, ADJ, and WM contributed to the study concept and design. KDT, Y-DIC, RT, JIR, OHF, MD, A Petersman, Y-PF, DMB, RAM, LCB, WK, JWK, PMR, DIC, TM, PAP, JAS, SLRK, MEK, GD, MW, KS, MM-N, EB, MPMdeM, NS, AGU, WM, and BMP performed data acquisition. Data analysis was performed by XG, JY, TK, SG, Y-PF, LRY, CKW-C, FG, DIC, JAB, LFB, MEK, KLW, ACM, PSdeV, AD, JEH, and Y-PF. The manuscript was drafted by CKW-C, ACM, and PSdeV. Critical revision of the manuscript included contributions from GDS, HG, FPH, JB, XG, TK, OHF, CJO’D, SG, MD, A Petersman, LRY, NP, WK, JWK, CKW-C, PMR, DIC, SJB, LFB, PAP, JAS, SLRK, MEK, GD, NLS, WT, ACM, PSdeV, EB, MPMdeM, AD, JEH, ADJ, CS, NS, AGU, BMcK, WM, and BMP. Funding for the manuscript was provided by JIR, OHF, CJO’D, A Petersman, DMB, RAM, LCB, WK, JWK, DIC, SLRK, A Peters, KS, MPMdeM, AD, ADJ, WM, and AGU

## Supporting Information Legends

Supplemental Methods and Tables.doc

File containing the Supplemental Online Methods (including cohort specific information) as well as the Supplemental Tables (1-5)

